# AMPK is dispensable for physiological podocyte and glomerular functions but prevents glomerular fibrosis in experimental diabetes

**DOI:** 10.1101/2025.04.07.647592

**Authors:** Swayam Prakash Srivastava, Olivia Kopasz-Gemmen, Abhiram Kunamneni, Aaron Thurnman, Eden Ozukan, Vinamra Swaroop, Shota Yoshida, Sungki Hong, Ken Inoki

## Abstract

AMP-activated protein kinase (AMPK) has been postulated to be crucial in regulating various renal physiology and pathophysiology processes, including energy metabolism, ion and water transport, inflammation, and hypertrophy. However, the specific roles of AMPK in the podocyte, a cell critical for maintaining glomerular filtration, have not been fully explored using genetic model animals. In this study, we generated mice lacking both AMPK α1 and α2 catalytic subunits in glomerular podocytes (pmut). Our findings revealed that, surprisingly, AMPK is dispensable for normal podocyte function. These knockout mice could live as long as their wild-type littermates without showing any pathological alterations in their glomeruli or glomerular function at two years of age. However, under type 1 diabetic conditions, the diabetic pmut mice exhibited increased lipid and collagen accumulation and an elevated expression of mesenchymal proteins in their glomeruli. They also showed more significant albuminuria compared to control diabetic mice. Under high glucose culture conditions, glomeruli isolated from pmut mice demonstrated a reduced expression of mitochondrial genes (e.g., Ndufv2) and increased leakage of mitochondrial components. Additionally, there was heightened expression of genes associated with nucleotide sensing and pro-inflammatory pathways (including mb21d2, IL-1 beta, and NF-kB). These observations suggest that while AMPK is not necessary for podocyte function in healthy kidneys, it is crucial for preventing glomerular fibrosis resulting from lipotoxicity and inflammation under diabetic conditions.

## Introduction

AMP-activated protein kinase (AMPK) is an evolutionarily conserved serine/threonine protein kinase, of which kinase activity is stimulated by various cellular stresses. Upon activation, AMPK stimulates cellular catabolic processes, including beta-oxidation, glycolysis, and autophagy, to restore cellular energy levels. AMPK forms a hetero-trimeric protein complex in which the α subunits contain a kinase domain while β (β1 and β2) and γ (γ1-3) subunits function as scaffolds and regulatory components in the complex ^1,2^. The mammalian AMPK α subunit has two isoforms (α1 and α2), and the α1 subunit is ubiquitously expressed in all the tissues, but the α2 subunit is predominantly expressed in skeletal and cardiac muscles^2,3^. In response to energy limitation, increased cellular AMP or ADP directly interacts with the gamma subunit of AMPK and supports the kinase activity of the alpha subunit^4,5^. Moreover, AMPK can also sense cellular glucose levels independent of cellular adenine nucleotide levels^6^. Recent studies demonstrated that aldolase, a key enzyme in the glycolysis pathway that catalyzes fructose-1,6-bisphosphate (FBP), acts as a glucose metabolite-sensing protein for AMPK activation. Under glucose-limited conditions, FBP-unoccupied aldolase stimulates the recruitment of LKB1, a key upstream of AMPK, to the lysosome, where the LKB1 directly phosphorylates and activates AMPK^7^.

Given the critical roles of AMPK in cellular catabolic processes, it has been postulated that the reduction of AMPK activity contributes to the development of metabolic disorders such as diabetes, obesity, and their complications. Diabetic kidney disease (DKD) is the leading cause of end-stage renal disease, which affects almost one-third of diabetic patients worldwide^8–10^. Loss of glomerular function and glomerular fibrosis is the final consequence of DKD^8^. While a plethora of pathogenic factors leading to renal cell dysfunctions have been proposed to develop DKD^8,11–21^, dysfunctions of the podocyte, a highly differentiated glomerular epithelial cell, play a pivotal role in the development of DKD. Furthermore, recent studies proposed that the reduction of AMPK activity in podocytes under diabetic conditions contributes to the development of DKD^22–25^. In support of the above, compounds (e.g., metformin, resveratrol, AICAR, and other compounds activating AMPK through the beta subunit) that indirectly or directly activate AMPK activity mitigate podocyte injuries and other renal cell dysfunctions under diabetes-mimetic cell culture system or in diabetic model animals^26–30^. However, most observations showing reduced podocyte damage or the renoprotective effect of AMPK come from cell-culture experiments and mice treated with AMPK activators, which may have off-target actions and glucose-lowering effects. Thus, whether a reduction in AMPK activity in podocytes has an autonomous impact on the development of podocyte and glomerular dysfunction remains largely elusive. To examine the role of AMPK in podocytes for glomerular function in vivo, we generated podocyte-specific AMPK α1/α2 double knockouts (pmut) mice, in which both α catalytic kinase subunits were ablated. Surprisingly, we found that AMPK is dispensable for podocyte growth, survival, and glomerular function under ad-lib fed conditions, as pmut mice can live as long as wild-type mice without manifesting any pathological alteration in their kidneys. However, under type 1 diabetic conditions, pmut mice display the elevation of collagen synthesis in podocytes and of the endothelial-to-mesenchymal transition in glomerular endothelial cells, and the cumulative effects lead to more glomerular fibrogenesis than wild-type diabetic mice. These observations indicate that under normal conditions, AMPK has little role in maintaining podocyte functions, or the key roles of AMPK in supporting podocyte functions can be sufficiently compensated by other kinases. However, under diabetic conditions, the remaining AMPK activity within the podocytes in the wild-type mice does play a renoprotective role in mitigating podocyte phenotypic changes and related glomerular endothelial activation.

## Material and Method

### Antibodies and Reagents

Rabbit anti-pS6 (Cat: 5364), Rabbit CPT1a (Cat: 12252), Rabbit LC3A (Cat: 4599) and rabbit PKM2 (Cat: 4053) were purchased from Cell Signaling Technology. Anti-mouse WT-1 (Cat: 05-753) was purchased from EMD. Rabbit p-ULK-1 (Cat: 80218-1-RR), Rabbit anti-PFKFB3 (Cat: 13763-1-AP), Rabbit anti-Fibronectin (Cat: 15613-1-AP), and p-IKB (Cat: 82349-1-RR) were purchased from Proteintech. Goat anti-CD31 (Cat: AF3628) was purchased from the R&D System. Anti-TFAM (Cat: PA5-29571), rabbit anti-Vimentin (Cat: MA5-35320), and Rabbit anti-Collagen I (Cat: PA5-95137) were obtained from Invitrogen. Fluorescence-, Alexa Fluor 647–, and rhodamine-conjugated secondary antibodies were obtained from Life Technology. Streptozotocin was purchased from Sigma.

### Animal Experiments

For the podocyte-specific loss of function of AMPK, we bred the *Ampk α1 ^fl/fl^α2 ^fl/fl^* mice with *podocin-Cre* mice to generate mice with deletion of all catalytic subunits of AMPK, specifically in podocytes (pmut mice). All mice were on the C57BL/6 background. We analyzed both males and females in the experiments on aged mice. Given the worsened diabetic kidney disease phenotypes observed in male compared with female mice, male mice were used in the diabetic mice experiments. Briefly, type 1 diabetes was induced in 8 weeks of age of *Ampk α1 ^fl/fl^α2 ^fl/fl^*, *podocin-Cre* (pmut) mice and control littermates, *Ampk α1 ^fl/fl^α2 ^fl/fl^*, (WT) mice with five consecutive intraperitoneal doses of Streptozotocin (STZ: 50 mg/kg in 10 mmol/L citrate buffer (pH 4.5)), and the mice were monitored over 24 weeks^31–34^. Urine samples for albumin and creatinine levels were collected using metabolic cages at the indicated points (8, 12, and 20 weeks after STZ treatment). Before sacrifice, mice were weighed, and blood glucose levels were measured using glucose strips. All animal experiments were approved by the Institutional Animal Care and Research Advisory Committee at the University of Michigan.

### *In vivo* Physiologic Studies

Urinary albumin and creatinine concentrations were determined using a mouse-specific Albuwell M and Creatinine Companion kit (Exocell). Glomerular tuft area, PAS-positive mesangial area, and other pixel densities obtained by immunohistochemical experiments were measured using ImageJ software.

### Isolation of Glomeruli

The podocyte isolation was performed as previously described^35^. Adult C57BL/6 mice at 8-14 wk of age were used in this study. For glomerular isolation with the differential adhesion method, a mouse was euthanized by an overdose of Isoflurane. The two kidneys from one mouse were harvested, and the medulla was removed carefully. The renal cortexes were minced into tiny particles by chopping with a razor blade in 0.5 ml of HBSS and subjected to digestion with collagenase type V at 37°C for 15-20 min with pipetting at 5-minute intervals. The digestion was stopped by adding 4 ml of DMEM supplemented with 10% FBS. The digested tissue was spun down at 290 g for 1 min at 4°C and resuspended in 5 ml of HBSS. The resulting mixture was transferred onto a prewetted 100 µM plastic cell strainer. The strainer was washed with 5 ml of fresh cold HBSS. All filtrate was collected and washed through a prewetted 75 µM plastic cell strainer with 35 ml of fresh cold HBSS. The filtrate (50 ml) was collected and washed again through another prewetted 40 µM plastic cell strainer with the fresh cold HBSS. The retained glomerular and tubular fragments on the top of the 40 µM strainer were rinsed into a clean cell culture dish. After 12 min of settling, large, fragmented tubules adhered to the bottom of the dish while leaving most glomeruli and small fragmented tubules floating in the supernatant. With gently swirling the culture dish, the glomeruli-enriched supernatant was collected and transferred with a plastic Pasteur pipette onto a new prewetted 40 µM strainer. After a wash with 25 ml of fresh cold HBSS to remove the small fragments of tubules, the retained glomeruli on the top of the 40 µM strainer were rinsed into another clean cell culture dish for the second adhesion to remove the residual large fragments of tubules. The supernatant containing highly purified glomeruli was collected and centrifuged at 290 g at 4°C for 5 min in a centrifuge equipped with a swing-out rotor.

### Kidney Histology

Sirius red, periodic acid-Schiff, and Masson-Trichrome staining were performed by the University of Michigan pathology Core and visualized using an Aperio imaging system. Masson-Trichrome-stained sections were evaluated using ImageJ software, and the fibrotic areas were estimated. For Sirius red staining, deparaffinized sections were incubated with picrosirius red solution for 1 hour at room temperature. The slides were washed twice with acetic acid solution for 30 seconds per wash. Then, the slides were dehydrated in absolute alcohol 3 times. The slides were cleared in xylene and mounted with a synthetic resin. For each mouse, images of 8 different fields of view were evaluated at ×40 magnification, and 15 to 20 stained glomeruli from each mouse were analyzed. Masson-Trichrome stain, a relative area of fibrosis, and Sirius red relative collagen deposition were analyzed to calculate glomerular fibrosis and glomerular collagen deposition, respectively. Glomerular surface area was calculated using traced glomeruli and ImageJ algorithms.

### Immunofluorescence

Paraffin-embedded kidney sections (5 μm) were used for immunofluorescence staining; double-positive labeling with pS6, LC3, Vimentin, Desmin, Collagen I, Biodipy, CPT1a, PFKFB3, PKM2, TFAM, pIKB, p65, or pATG1 with WT1 for podocytes; and Vimentin, Fibronectin, CPT1a, PFKFB3, or PKM2 and CD31 for endothelial were performed.

### Image analysis and quantification

Data were analyzed using ImageJ or Fiji ImageJ version 1.54p. Immunofluorescence intensity was determined by ImageJ by splitting the color channel, and the intensity of the signal was divided by glomerular area (intensity/µm^2^). The number of LC3 puncta in the glomerulus was determined by Fiji Image J. Briefly, glomerular regions of the fluorescence images were cropped, their color channels were split, and the background was reduced. The ratio (the number of LC3 puncta/glomerular area) was automatically determined using H-watershed (hmin=7.0, thresh=18, peakflooding=35, allowsplitting=true) and monitored the number of particles (size=0-50: less than 8 µm^2^).

### Oil red O staining

Briefly, 5 μm kidney sections were dried at room temperature for 15 min and fixed in prechilled acetone solution for 10 min. Slides were washed three times with PBS for 5 min and lastly rinsed in 60% isopropanol for 5 min. Lipids were stained by incubating slides in fresh Oil Red O working solution for 60 min at room temperature and rinsed in 60% isopropanol for 5 s. Slides were washed three times with PBS and 5 min with water and counterstained with hematoxylin. Last, slides were washed in 70% ethanol, mounted with mounting media, and immediately imaged.

### EndMT detection

Paraffin-embedded kidney sections (5 μm) were used for the detection of EndMT. Cells undergoing EndMT were detected by double-positive labeling for CD31 and Vimentin and CD31 and Fibronectin. Sections were analyzed and quantified by fluorescence microscopy.

### Transmission electron microscopy

The sample preparations and electron microscopic analyses were performed at the Biomedical Research Core at the University of Michigan. For sample preparation, mice were anesthetized and perfused with Sorensen phosphate buffer containing 4% PFA and 2.5% glutaraldehyde, and kidneys were isolated. The processed samples were analyzed by Philips CM100 TEM or AMRAY 1910 field emission scanning electron microscope. ImageJ was used to quantitatively analyze electron micrographs. Foot processes were measured from at least 50 μm of GBM for each mouse. Podocyte foot processes and GBM thickness were analyzed by ImageJ. Images were blinded by assigning integer numbers before evaluation by someone other than the scorer.

### mRNA isolation and Quantitative Polymerase Chain Reaction

Total RNA was isolated using the standard Trizol protocol. RNA was reverse transcribed using the iScript cDNA Synthesis kit (Bio-Rad), and quantitative polymerase chain reaction was performed on a Bio-Rad C1000 Touch thermal cycler using the resultant cDNA, quantitative polymerase chain reaction master mix, and gene-specific primers. The primers used are given in **Table S1**. Gene expression was normalized to the housekeeping gene actin and is presented as fold change. All primers were synthesized by the IDT.

### Statistical analysis

All values are expressed as mean ± SEM and analyzed using GraphPad Prism 7 (GraphPad Software Inc., La Jolla, CA). The significance of differences was determined using an unpaired two-tailed Student’s t-test or one-way ANOVA. p values of less than 0.05 were considered statistically significant: *p<0.05, **p<0.01, ***p<0.001, ****p<0.0001

## Results

### AMPK is dispensable for the maintenance of podocyte and glomerular function in the normal development and aging stages

To test the role of AMPK in podocytes, we generated podocyte-specific AMPK α1/α2 double knockout mutant mice (pmut) by crossing B6 background *Ampk α1 ^fl/fl^α2 ^fl/fl^* mice with B6 background *podocin-Cre* mice **(Figure 1A)**. The isolated glomeruli of the pmut mice had significantly suppressed levels of both AMPKα1 and AMPKα2 mRNAs when compared to the WT groups **(Figure S1A)**. The loss of the AMPK activity in podocytes was further analyzed by double staining of WT1 (podocyte marker) and phosphorylated ATG1 (S556), the site phosphorylated by AMPK. Consistent with the reduction of AMPKα1 and AMPKα2 mRNAs in the glomeruli, levels of pATG1 were diminished in the podocyte of pmut mice (**Figure S1B)**. Wild-type (WT) and the pmut mice were kept on free access to food and water for 2 years, and physiological parameters and kidney histology were monitored. The pmut mice (both male and female) were healthy as control mice throughout the monitoring period (2 years). There was no significant difference in body weight, kidney weight, and blood glucose at 2 years old of pmut mice and littermate controls **(Figure 1B, 1C, and S1C)**. Histological analyses also demonstrated that no pathological alterations of glomerular and tubular structures (Hematoxylin-Eosin: HE), fibrosis (Masson Trichrome: MTS), collagen deposition (Sirius red) in the pmut mice cortex tissues compared to control mice (**Figure 1D, 1E, and 1F)**. Periodic acid-Schiff (PAS) staining also indicated that there is no significant difference in the accumulation of carbohydrate molecules or glomerular surface area in the pmut mice compared to those in control mice (**Figure 1G)**. As expected, there were no apparent differences in urine albumin-to-creatinine ratios between the two groups **(Figure 1H)**.

**Figure 1.**
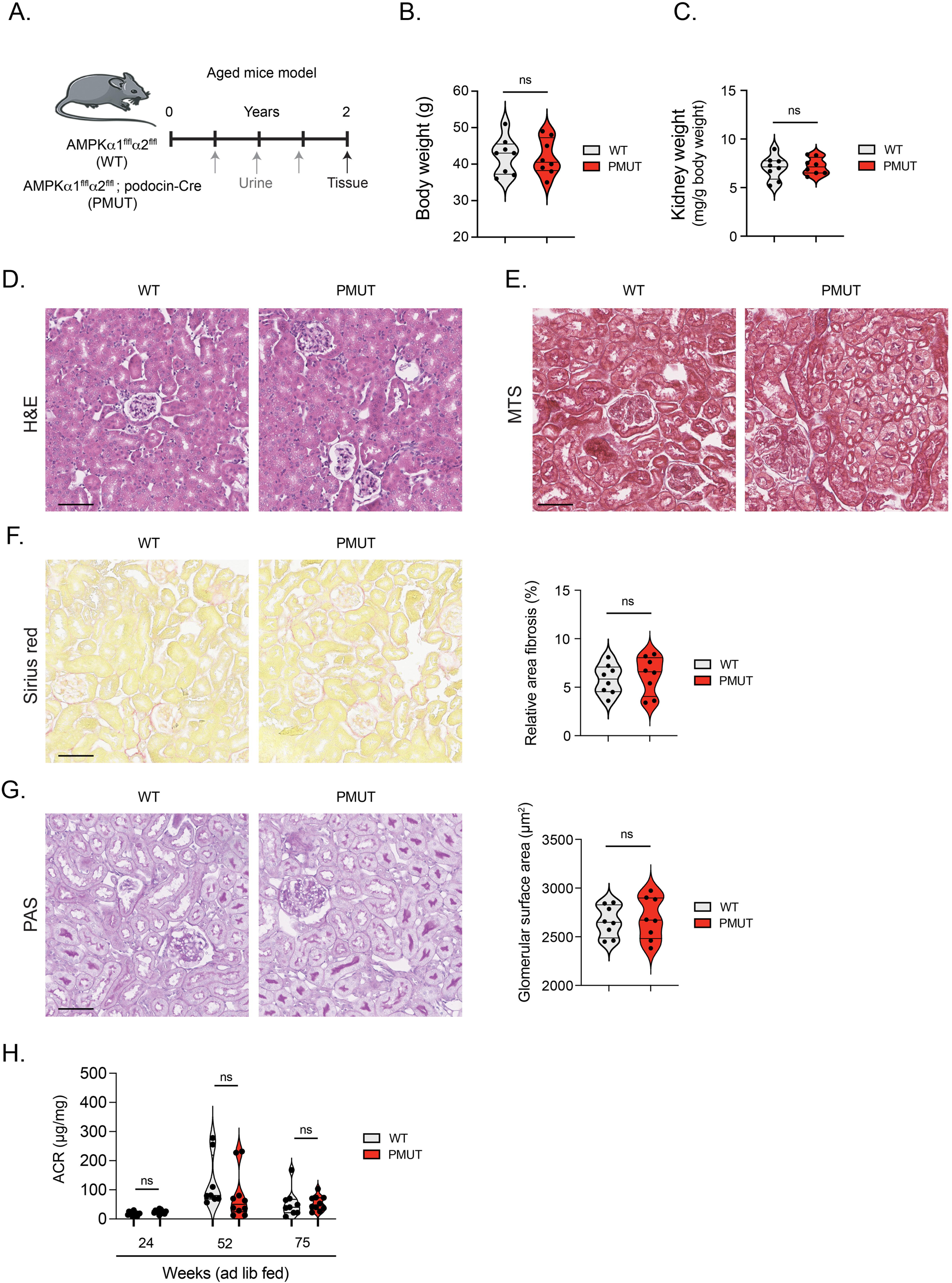
AMPK is dispensable for the maintenance of podocyte and glomerular function in the normal development and aging stages. **A.** A schematic diagram showing the experimental design analyzing the aged *Ampk α1 ^fl/fl^α2 ^fl/fl^*mice (WT) and *Ampk α1 ^fl/fl^α2 ^fl/fl^*, *podocin-Cre* mice (pmut). WT and pmut mice were kept on a normal chow diet and had free access to water. At the age of 2 years, mice were used for the analysis of histopathology. **B.** Measurement of body weight. Data are shown as mean±SEM. ns denotes no significance in the statistical analysis. N=8 was analyzed/group. **C.** Kidney weight/body weight was measured. Data are shown as mean±SEM. ns denotes no significance. N=8 was analyzed/group. **D, E.** Hematoxylin-eosin and Masson-Trichrome staining were shown. **F.** Sirius red staining was shown. The relative area of fibrosis in the glomeruli was quantified. Data are shown as mean±SEM. ns denotes no significance. N=8 was analyzed/group. **G.** Periodic Acid-Schiff (PAS) staining was shown. The glomerular surface area was calculated. The scale bar indicates 50 µm. Data are shown as mean±SEM. ns denotes no significance. N=8 was analyzed/group.**H.** Albumin-to-creatine ratio (ACR: µg/mg) was shown. 24-hour urine samples were collected at the indicated time points, and the albumin and creatinine concentrations were determined. Data are shown as mean±SEM. ns denotes no significance. N=8 was analyzed/group.

### AMPK in podocytes is dispensable to reduce cellular mTORC1 activity and maintain autophagy under normal development and aging conditions

AMPK is known to act as a key kinase that protects the vulnerability of cells in response to various metabolic stresses. However, the number of podocytes in the glomeruli of pmut mice was maintained and comparable to that in control mice at 2 years old (**Figure 2A**). While AMPK has been proposed to act as a key suppressor for mTORC1 activity in various cells, including podocytes, levels of S6 phosphorylation, a downstream of mTORC1-S6K1 activity, were not increased in the glomeruli of pmut mice compared to that in the glomeruli of wild-type mice (**Figure 2B**). While levels of S6 phosphorylation in the glomeruli are similar between female WT and pmut mice, it was somewhat lower in the male pmut mice compared to that in the WT mice, suggesting that AMPK did not suppress mTORC1 activity under ad-lib fed conditions. Recent studies proposed that, in contrast to the general paradigm, acute AMPK activation inhibits autophagy^36^. However, continuous AMPK activation still supports cellular autophagy by maintaining the expression of proteins involving autophagy induction. Furthermore, it has been reported that AMPK may play a central role in upregulating autophagy activity^37^in podocytes. While the number of LC3 puncta, which indicates an autophagosome, in the glomerulus of pmut mice was slightly decreased compared to that of WT mice, the podocytes of pmut mice still formed a number of autophagosomes, suggesting that basal autophagic activity was maintained in the podocyte lacking AMPK (**Figure 2C**). This idea is also supported by the previous observations that loss of autophagy in podocytes resulted in age-dependent late-onset glomerulosclerosis and proteinuria, whereas the pmut mice did not display such pathological phenotypes^38^. These observations suggest that AMPK may not be a key player in regulating mTORC1 and AMPK activity in podocytes and is dispensable for podocyte and glomerular function under ad-lib-fed conditions throughout the life of mice.

**Figure 2.**
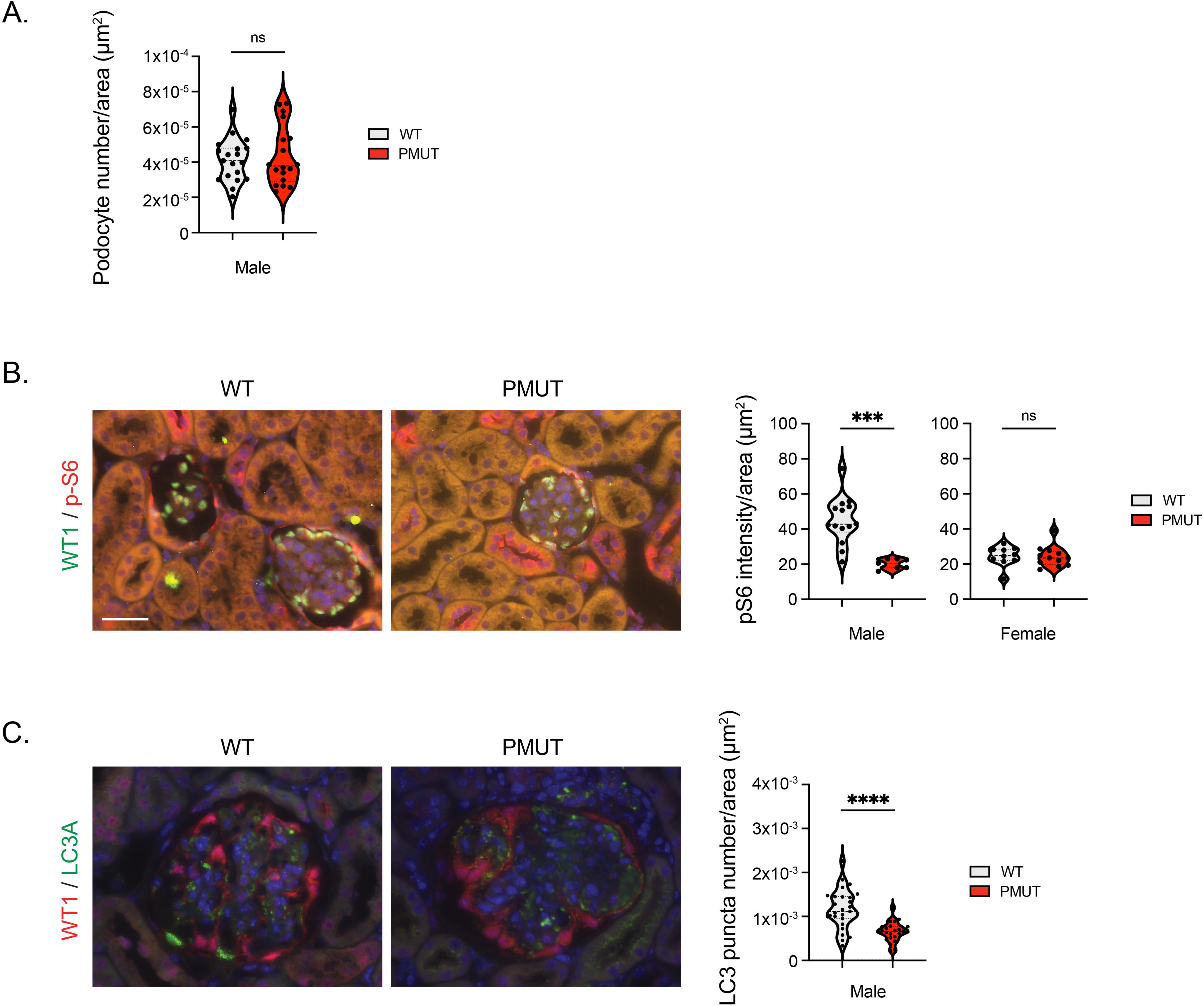
AMPK in podocytes is dispensable to reduce cellular mTORC1 activity and maintain autophagy under normal development and aging conditions. **A.** The number of WT1-positive podocytes per glomerulus (section) was determined in the indicated genotypes. Data are shown as mean±SEM. ns denotes no significance. N=18∼19 glomeruli from 5∼7 mice per group were analyzed. **B.** Levels of phosphorylated S6 were determined. Double staining for pS6 (S240/S244) and WT1 in the indicated genotypes was shown. The scale bar indicates 50 µm. The intensity of pS6 in the indicated glomeruli was quantified. Data are shown as mean±SEM. N=7∼14 images were analyzed per group. **C.** Double staining for WT1 and LC3 was shown. The number of LC3 puncta less than 8 µm^2^ in the indicated glomeruli was determined. Data are shown as mean±SEM. p****<0.0001. N=6/group were analyzed.

### AMPK loss in podocytes leads to enhanced glomerular fibrosis in a mouse model of type 1 diabetes

To investigate the role of AMPK in podocytes in the regulation of glomerular function under diabetic conditions, we induced diabetes in WT and pmut mice with Streptozotocin. Five low doses of Streptozotocin were injected intraperitoneally, and their glomerular function and histological alterations were monitored (**Figure 3A**). Both WT and pmut diabetic animals showed similar levels of hyperglycemia, body weight loss, and higher kidney weights compared to non-diabetic animals (**Figure 3B**, 3C**, and 3D**). As evidenced by transmission electron microscopy, glomerular ultrastructure showed no significant differences between non-diabetic WT and pmut mice. Both diabetic control and pmut mice displayed foot process effacement and GBM thickening (**Figures 3F and 3G**). While levels of GBM thickening in diabetic WT and pmut mice were similar, the foot process effacement of pmut podocytes was slightly more severe than that of control podocytes under diabetic conditions (**Figure 3F and 3G**). Furthermore, the histological analysis with PAS staining showed a slightly increased glomerular size and PAS-positive area in the glomeruli of pmut diabetic mice compared to those of WT diabetic mice (**Figure 3H, 3I, and 3J**). Consistent with these observations, pmut diabetic mice displayed higher levels of urine albumin-to-creatinine ratios (ACR) compared to diabetic WT mice (**Figure 3K**). These observations suggest that AMPK does play an important role in preventing podocyte dysfunctions in these diabetic model animals.

**Figure 3.**
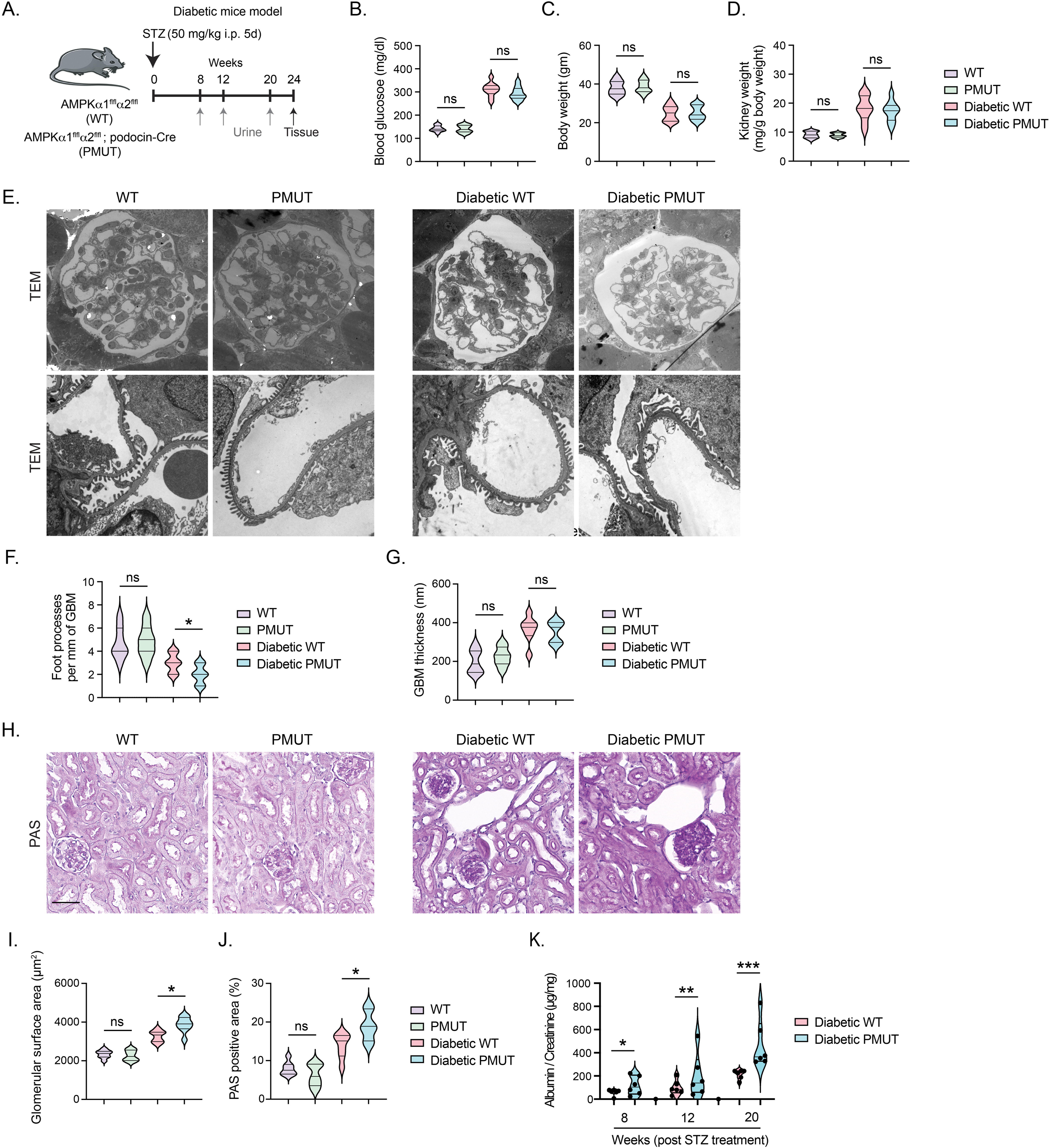
AMPK loss in podocytes leads to enhanced glomerular fibrosis in a mouse model of type 1 diabetes. **A.** A schematic diagram showing the experimental design analyzing the diabetic *Ampk α1 ^fl/fl^α2 ^fl/fl^*mice (WT) and *Ampk α1 ^fl/fl^α2 ^fl/fl^*, *podocin-Cre* mice (pmut). The indicated 8∼9 weeks old mice were subjected to the administration of five low doses (50 mg/kg body weight) STZ treatment. Diabetic mice were selected based on their blood glucose above 200 mg/dl, kept on chow food, and had free access to water for 24 weeks. 24-hour urine collection was performed at the indicated time points, and tissues were analyzed 24 weeks after the STZ treatment. At the end of the experiment, mice in each group were sacrificed and analyzed for kidney histology. **B, C, and D.** Blood glucose, body weight, and kidney weight/body weight were shown (24 weeks post STZ treatment). Data are shown as mean±SEM. N=10 was analyzed/group. **E**. Representative transmission electron microscopy images were shown. N = 3/group. **F, G.** Foot process effacement and glomerular basement membrane thickness were analyzed using the TEM images, and quantification was calculated using ImageJ software. Six independent images were analyzed. **H.** Periodic Acid-Schiff (PAS) staining was shown in the indicated group. **I, J.** Glomerular surface area was measured using the PAS images. The scale bar indicates 50 µm. Data are shown as mean±SEM. N=6 was analyzed/group. **K.** Albumin-to-creatine ratio (ACR) was determined by performing ELISA in diabetic WT and pmut mice. Data are shown as mean±SEM. N=6 was analyzed/group.

To investigate more pathological alterations in pmut diabetic mice, we perform Masson’s trichrome and Sirius red staining to evaluate levels of tissue fibrosis. These assays demonstrated an increased collagen deposition in mesangial and tubulointerstitial areas in both diabetic groups compared to non-diabetic groups. There was no difference in levels of interstitial fibrosis between the diabetic groups (**Figure 4A and 4B**). However, consistent with the observations in PAS staining (**Figure 3H and 3J)**, immunofluorescence studies revealed a higher expression of mesenchymal proteins, such as vimentin and desmin, and a higher level of collagen I deposition in the glomeruli of diabetic pmut mice compared to those of WT diabetic mice, indicative of higher glomerular fibrosis in diabetes pmut mice (**Figure 4C**).

**Figure 4.**
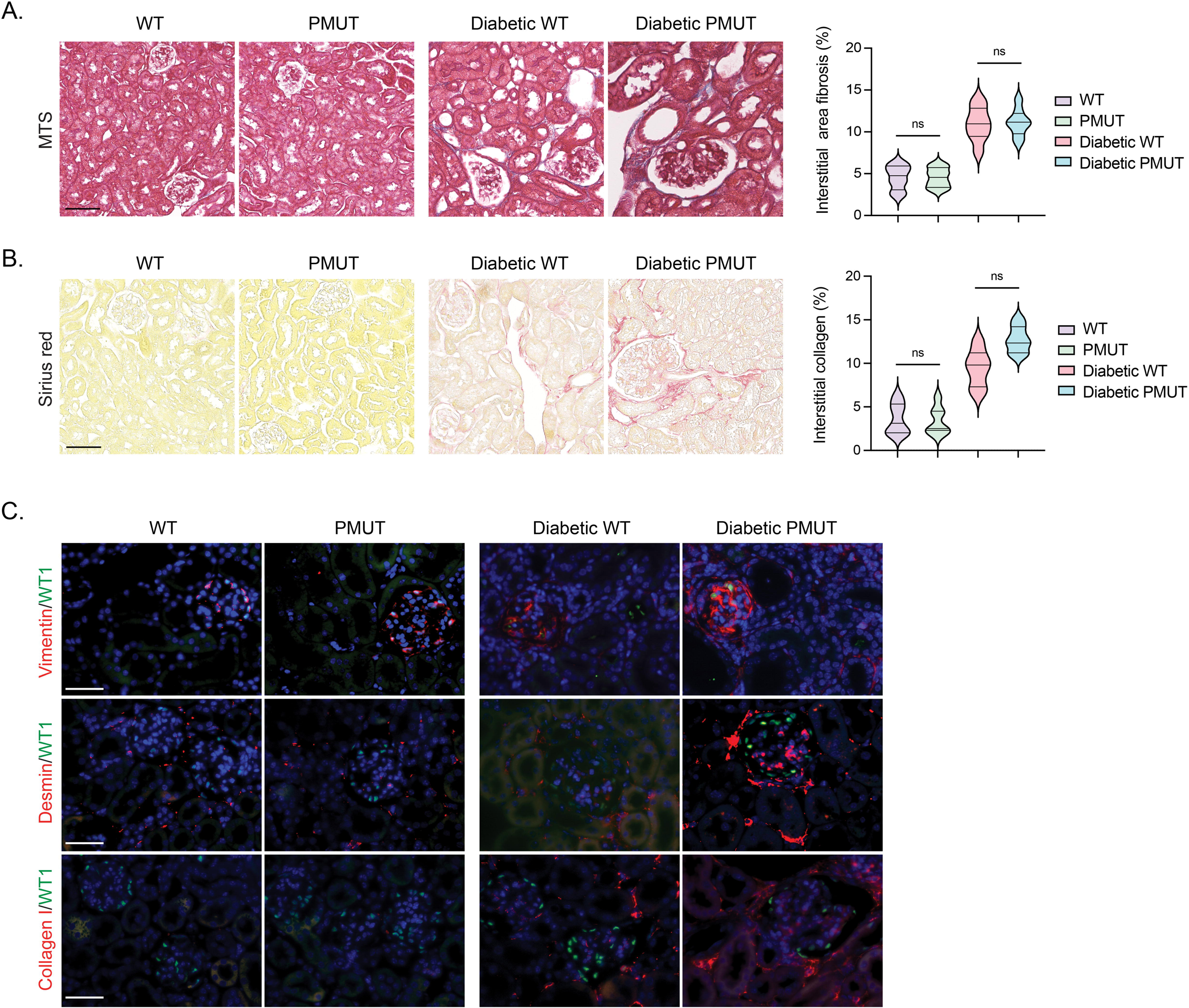
AMPK loss in podocytes leads to enhanced glomerular fibrosis in a mouse model of type 1 diabetes (continued). **A.** Masson Trichrome staining (MTS) was performed, and levels of fibrosis in the interstitial region were quantified. The scale bar indicates 50 µm. Data are shown as mean±SEM. N=10 was analyzed/group. **B.** Sirius red staining was performed, and levels of collagen deposition in the interstitial region were quantified. The scale bar indicates 50 µm. Data are shown as mean±SEM. N=10 was analyzed/group. **C.** Double immunofluorescence staining of vimentin, desmin, or collagen I with WT1 was shown. The scale bar indicates 50 μm. Representative images are shown. N=7 was analyzed/group.

### AMPK loss in podocytes leads to the alterations of lipid and glucose metabolisms in the mouse model of type 1 diabetes

Metabolic shifts play critical roles in the health and disease processes of kidney cells, and recent studies suggest that such alterations in fuel preference are associated with fibrogenesis in kidney cells^39–48^. AMPK is a known kinase that breaks down lipids to generate energy source for cell survival. We examined levels of lipid accumulation in kidney cells by performing Oil-red O and Bodipy assays. While there was no remarkable difference in lipid deposition in the kidney of non-diabetic WT and pmut mice (**Figure 5A** and **S1D**), overall, levels of Oil-red O-positive tissues were increased under diabetic conditions. Importantly, the glomeruli of diabetic pmut mice displayed a higher level of lipid deposition compared to the glomeruli of diabetic WT mice **(Figure 5A**). Immunofluorescent staining of Biodipy, which stains most of the neutral lipids, also revealed no remarkable difference in lipid deposition in the kidneys of non-diabetic WT and pmut mice. However, consistent with the results of Oil-red O staining, the diabetic pmut mice displayed higher levels of Biodipy-positive podocytes in the diabetic pmut mice compared to diabetic WT mice (**Figure 5B**). Moreover, levels of carnitine palmitoyltransferase 1a (CPT1a), a key enzyme in fatty acid oxidation (FAO), were largely reduced in diabetic kidney tissues in both genotypes (**Figure 5C**). Furthermore, levels of CPT1a expression in the podocytes of pmut diabetic mice were lower than that in WT diabetic mice (**Figure 5C)**, suggesting that the podocytes in diabetic pmut mice may have a more significant limitation in utilizing lipids and generating cellular energy from FAO. We also monitored levels of key enzymes in the glycolysis pathway. Under diabetic conditions, levels of 6-phosphofructo-2-kinase/fructose-2,6-biphosphatase 3 (PFKFB3) and pyruvate kinase muscle type 2 (PKM2) expression in podocytes were increased in both genotypes compared to those in the podocytes of non-diabetic mice. Interestingly, more prominent enhanced expression of these glycolytic enzymes was observed in the glomerular cells, including podocytes, of diabetic pmut mice (**Figure 5C**). While AMPK has been recognized as a key stimulator of glycolysis in response to metabolic stresses, these observations suggest that loss of AMPK in podocytes may lead to further enhancement of glycolysis to generate cellular fuel in the podocytes and other glomerular cells under type 1 diabetic conditions.

**Figure 5.**
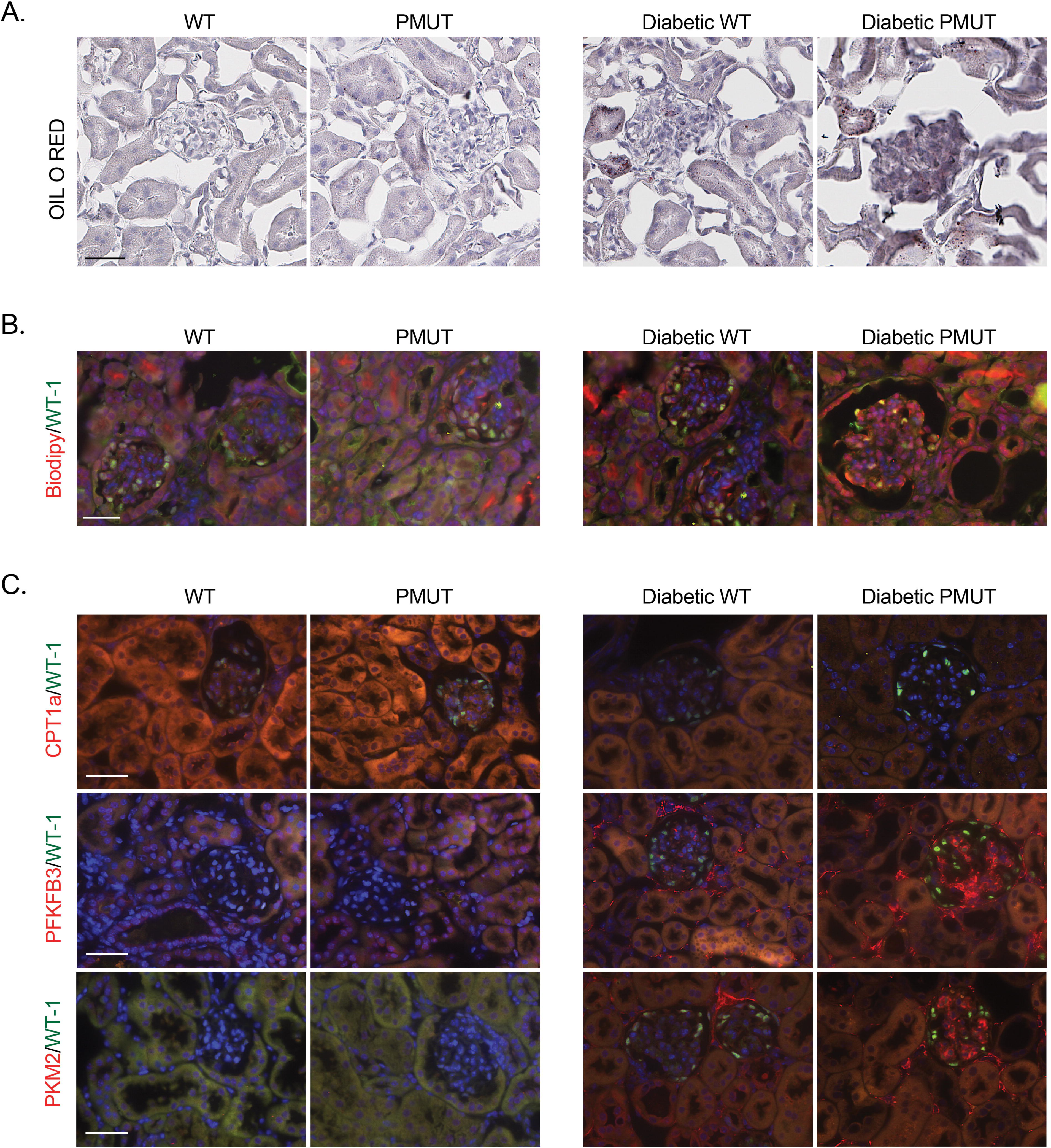
AMPK loss in podocytes leads to the alterations of lipid and glucose metabolisms in the mouse model of type 1 diabetes. **A.** Oil O Red staining was performed in non-diabetic and diabetic WT and pmut mice. Representative microphotographs are shown. The scale bar indicates 50 µm. N=6 was analyzed per group. **B.** Double immunofluorescence staining of Biodipy and WT1 was performed in the indicated genotypes. The scale bar indicates 50 µm. Representative images are shown. N=6 was analyzed/group. **C.** Double immunofluorescence staining of CPT1a, PFKFB3, or PKM2 with WT-1 was performed. The scale bar indicates 50 µm. Representative images are shown. N=6 was analyzed/group.

### AMPK loss in podocytes facilitates endothelial-to-mesenchymal transition in the mouse model of type 1 diabetes

Endothelial-to-mesenchymal transition (EndMT) plays a key role in the development of glomerular fibrosis ^21,39,49–56^. Since we observed that diabetic pmut mice displayed more glomerular fibrosis than diabetic control mice, we investigated whether loss of AMPK in podocytes facilitates EndMT in the glomerular endothelial cells under diabetic conditions. Double immunofluorescence staining with CD31, an endothelial marker, and vimentin or fibronectin revealed that higher levels of vimentin or fibronectin expression in the CD31-positive endothelial cells in the glomeruli of diabetic mice compared to those of non-diabetic mice. Importantly, this effect was more prominent in the glomeruli of diabetic pmut mice when compared to the glomeruli of the diabetic control mice (**Figure 6A**), indicating that loss of AMPK in podocytes facilitates EndMT of glomerular endothelial cells under diabetic conditions. Impairment of normal lipid or glucose metabolism has been linked to the EndMT phenotype in the endothelial cells ^39,40,57^. Therefore, we also analyzed levels of CPT1a and glycolytic enzymes in the endothelial cells. Consistently, overall levels of CPTa expression were reduced in the kidney tissues of diabetic groups compared to non-diabetic counterparts (**Figure 6B**). There was a consistent trend that levels of CPT1a were more reduced in the glomeruli and CD31-positive endothelial cells of diabetic pmut mice when compared to the diabetic control mice (**Figure 6B**). In contrast, levels of PFKFB3 and PKM2 expression were again enhanced in the glomeruli of diabetic mice than non-diabetic mice, and their expression in CD31-positive endothelial cells was much higher in the diabetic pmut mice compared to those in diabetic control mice (**Figure 6B**). These observations indicate that loss of AMPK in the podocytes also leads to similar metabolic derangements in the glomerular endothelial cells and facilitates EndMT under diabetic conditions.

**Figure 6.**
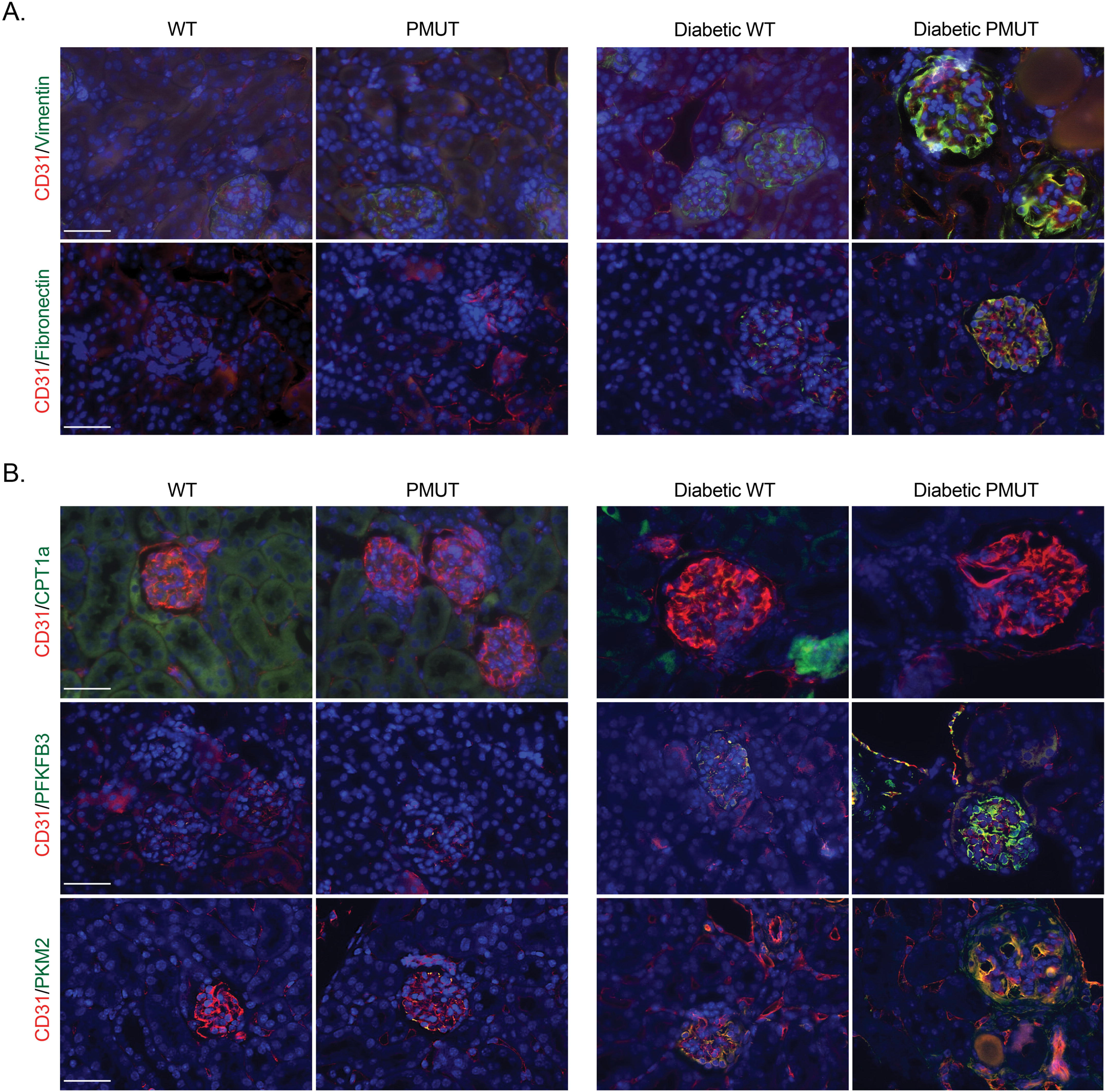
AMPK loss in podocytes facilitates endothelial-to-mesenchymal transition in the mouse model of type 1 diabetes. **A.** Endothelial-to-mesenchymal transition was determined by double immunofluorescence staining of vimentin or fibronectin with CD31. Representative immunofluorescence images are shown here. The scale bar indicates 50 µm. N=6 was analyzed/group. **B.** Double immunofluorescence staining of CPT1a, PFKFB3, or PKM2 with CD31 was performed. The scale bar indicates 50 µm. Representative images are shown. N=6 was analyzed/group.

### Loss of AMPK in podocytes causes c-GAS-STING-mediated inflammation

Metabolic reprogramming, such as defective normal lipid metabolism and associated lipotoxicity, is detrimental to the cells and cell organelles, including mitochondria ^58,59^. Damaged mitochondria often release mitochondrial DNA into the cytoplasm, triggering the activation of cytosolic GMP-AMP synthase (cGAS) and stimulator of interferon genes (STING) pathway, a critical innate immune response^60,61^. The activation of this pathway enhances the transcription of pro-inflammatory genes ^58,60^. To investigate mitochondrial integrity and expression in diabetic pmut mice, we monitored levels of mitochondrial transcription factor A (TFAM). Overall, the levels of TFAM expression were decreased in the kidney tissues of both genotypes under diabetic conditions. Importantly, the glomeruli and podocytes of diabetic pmut mice displayed reduced levels of TFAM compared to those of control diabetic mice **(Figure 7A**). To test whether these observations can be recapitulated in the in vitro system, glomeruli were isolated from the kidney cortex tissues of pmut and control mice (10-week-olds) and cultured in high glucose media. Consistently, levels of *Tfam* and *Ndufv2* mRNAs were lower in the glomeruli of pmut mice compared to those of control mice **(Figure 7B)**. To test if loss of AMPK leads to reduced mitochondria integrity, we monitored levels of *mt-Co1* and *mt-Cyb* transcripts in the cytosolic fractions of cultured glomeruli. Levels of cytosolic *mt-Co1* and *mt-Cyb* transcripts were increased in the glomeruli of pmut mice compared to those in control mice, suggestive of more leakage of mitochondrial components in the pmut glomerular cells under high glucose culture conditions **(Figure 7C)**. Furthermore, levels of the pro-inflammatory gene expression, such as *IL-1*β, *IL-6*, and *NF-*κ*B1,* were significantly higher in the glomeruli from pmut mice than those from control mice under high-glucose culture conditions **(Figure 7D)**. The gene expression of *Mb21d2 (*cGAS) and *Tmem173* (STING) is upregulated by various transcriptional factors, including NF-κB and STAT1^58,60,62^. Consistently, levels of *Mb21d2 and Tmem173* were higher in the glomeruli of pmut mice than those in control mice (**Figure 7E**), suggesting that loss of AMPK predisposes to augment mitochondrial-derived inflammation through the cGAS-STING pathway under high glucose conditions. We also observed that the isolated glomeruli from the pmut mice expressed a higher level of lactate dehydrogenase (LDH) mRNA expression, whereas lower levels of pyruvate dehydrogenase (PDH) mRNA expression compared to control glomeruli under high glucose culture conditions (**Figure 7F**), suggesting that loss of AMPK in podocytes may also lead to a metabolic shift in the glycolytic pathway to produce lactate, which has been found to accelerate the fibrotic process in many cell types^26,63–67^.

**Figure 7.**
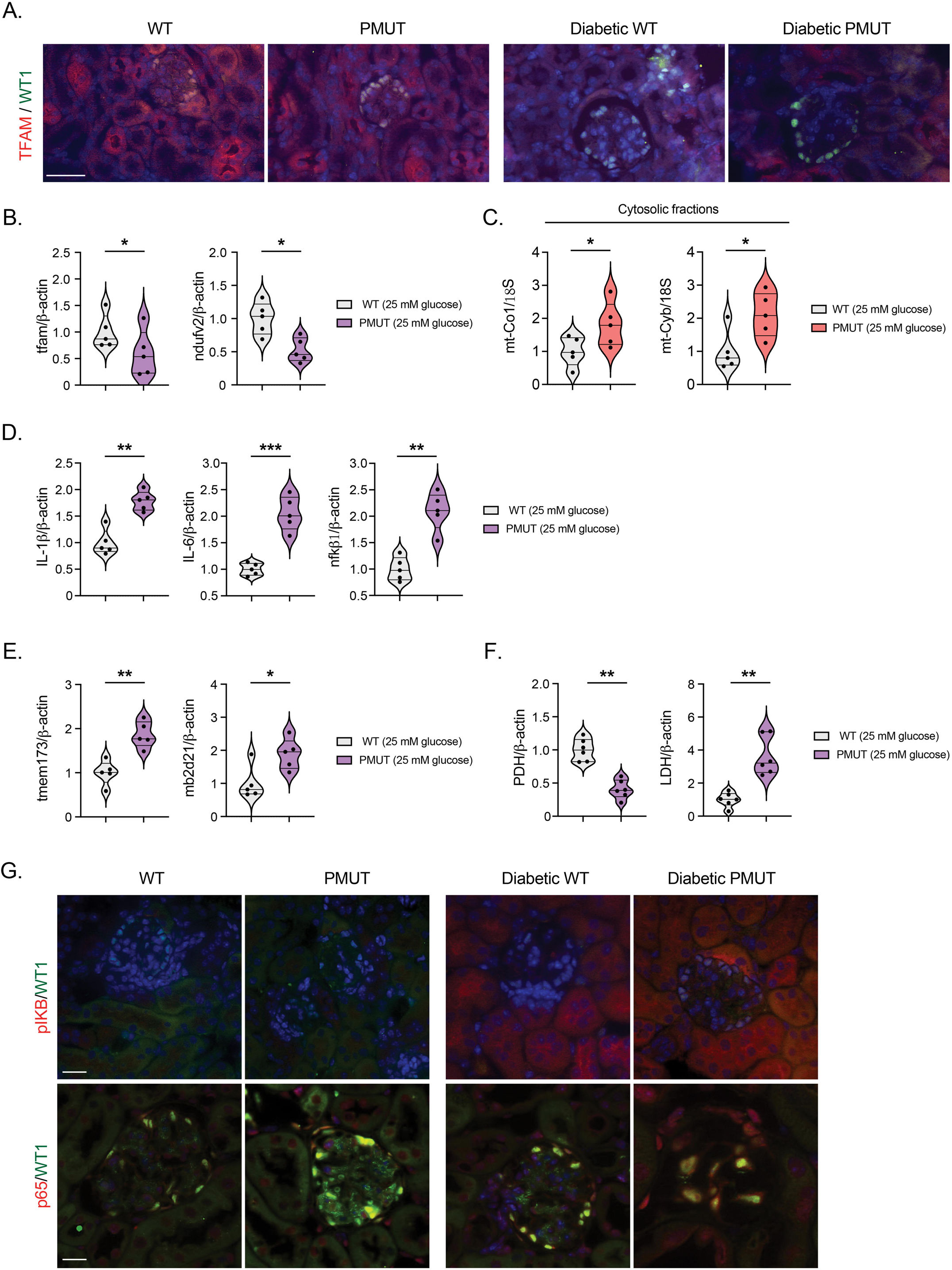
Loss of AMPK in podocytes causes c-GAS-STING-mediated inflammation. **A.** Double immunofluorescence staining of TFAM and WT-1 was shown in the non-diabetic and diabetic WT and pmut mice. The scale bar indicates 50 µm. Representative images are shown. N=6 was analyzed/group. **B.** Quantitative qPCR analyses of tfam and ndufv2 transcripts in the indicated glomeruli were shown. mRNAs were extracted from the purified glomeruli from WT or pmut mice under high-glucose (25 mM) culture conditions (for 96 hours). Data are shown as mean±SEM. N=6, *p<0.05. **C.** Quantitative qPCR analyses of mt-Co1 and mt-Cyb transcripts in the cytosolic fractions of high-glucose (25 mM) treated cultured glomeruli from WT and Pmut mice. 18S was used as an internal control. Data are shown as mean±SEM. N=6, *p<0.05. **D, E.** Quantitative qPCR analyses of il-1β, il-6, nfkb1, tmem173, and mb2d21 transcripts in the indicated glomeruli cultured in high-glucose (25 mM) media. Data are shown as mean±SEM. N=6, *p <0.05, **p <0.001, ***p <0.0001. **F.** Quantitative qPCR analyses of phd and ldh transcripts in the indicated glomeruli cultured in high-glucose (25 mM) media. Data are shown as mean±SEM. N=6, **p <0.001. **G.** Double immunofluorescence staining of p-IKβ or p65 with WT1 was shown in non-diabetic and diabetic WT and pmut mice. The scale bar indicates 50 µm. Representative images are shown. N=6 was analyzed/group.

Finally, immunofluorescent staining demonstrated that levels of phospho-IκB were generally higher in the diabetic kidney tissues, including glomerulus and tubular cells in both groups and the intensity was higher in the glomeruli of diabetic pmut mice when compared to that in control diabetic mice (**Figure 7G)**. Accordingly, nuclear p-65 expression accumulated more in the glomerular cells, especially in podocytes, of diabetic pmut mice compared to those of control diabetic mice (**Figure 7G)**. These observations suggest that loss of AMPK leads to lipid accumulation, mitochondrial damage, and the activation of the cGAS-STING pathway, resulting in inflammation in type 1 diabetic mice.

## Discussion

Podocytes are highly differentiated epithelial cells. While they lack the capability of replication, they are metabolically active to maintain the functions of the filtration barrier as well as the glomerular basement membrane by operating high levels of both anabolic (e.g., protein synthesis) and catabolic (e.g., autophagy) processes. AMPK is known to function as a central regulator that stimulates catabolic processes in various cells, including podocytes. Many previous reports indicated that reduction of AMPK activity plays a key role in developing DKD based on the two categories of biological observations. Firstly, AMPK activity, which is inhibited by not only cellular energy (e.g., ATP) but also glucose metabolites (i.e., FBP), is generally reduced in the diabetic kidney in which glomerular cells are exposed to high levels of glucose and nutrients. Second, diabetic mice treated with compounds that directly or indirectly enhance AMPK activity display some renoprotective effects against DKD development in several animal models^26,27,67–74^. Furthermore, these AMPK activators reduce the vulnerability of podocytes caused by metabolic stresses related to diabetes (e.g., high concentrations of glucose) in the cell culture system. In addition to these observations, aberrant mTORC1 activation in podocytes has been observed in diabetic animals and patients and causes pathological podocyte hypertrophy and detachment from the GBM and is thought to contribute to the development of DKD^75–77^. Importantly, AMPK acts as a key suppressor of mTORC1 and may enhance autophagy (at least in long-term AMPK activation) in various types of cells^36,78–80^, including cultured podocytes, supporting the idea that reduction of AMPK activity in podocytes under diabetic conditions may underlie in the pathogenesis of podocyte dysfunction and DKD. If this idea is correct, the ablation of AMPK in podocytes likely leads to podocyte vulnerability and injury autonomously and may manifest DKD-like phenotypes as seen in the model where mTORC1 activity is genetically activated specifically in podocytes^76,81^. However, no report has been reported on the glomerular phenotypes of podocyte-specific AMPK knockout mice, in which both α1 and α2 catalytic subunits are ablated. Hence, we report here that, in contrast to our prediction, AMPK is dispensable for podocyte development and its function in the normal developmental stage and aging processes as the pmut KO mice do not exhibit any pathological glomerular alterations at two years old compared to their control littermates. These observations were somewhat unexpected given the reported AMPK’s various key roles in maintaining energy homeostasis, metabolic processes, and cytoskeletal organizations. Intriguingly, levels of mTORC1 activity were not significantly increased in the podocytes of pmut mice compared to that in control mice under normal ad-lib fed conditions. Furthermore, there is no obvious impairment of autophagy in the podocytes of pmut mice. These observations suggest that AMPK is likely inactive in the podocyte under normal developmental conditions or that other pathways could completely compensate for the functions of AMPK in maintaining podocytes. It has been reported that the activity of autophagy in podocytes was not inhibited by canonical mTORC1 activity, as autophagy is still active in the podocytes lacking the functional TSC complex, in which canonical mTORC1 is highly active^82^. The study proposed that AMPK may play a key role in maintaining autophagy activity in the podocyte. However, our study suggests that under normal physiological conditions, AMPK is also dispensable for maintaining autophagy activity in podocytes. While AMPK activation provides beneficial effects on stressed podocytes, the podocytes in mice can maintain their healthy growth and essential filtration and other glomerular functions without AMPK.

It has been postulated that AMPK activity is reduced in many tissues, including kidney cells (e.g., tubular cells and podocytes), in both type1 and type2 diabetic animals^26,67,83–91^. Our genetic studies suggested that the remaining reduced AMPK activity plays an important role in mitigating the progression of podocyte and glomerular dysfunction, especially under type 1 diabetic conditions. Mechanistically, we demonstrated that the remaining AMPK activity in podocytes is critical to mitigate lipid accumulation, mitochondria damage, and inflammation in podocytes and the phenotypic changes of glomerular endothelial cells. Consistent with the well-known role of AMPK in stimulating lipid catabolism, loss of AMPK led to the accumulation of lipids in the podocytes of type 1 diabetes, under which podocytes may uptake more FFAs from the filtration area. The reduction of CPT1a expression further exaggerates cytosolic lipid retention and likely causes lipotoxicity to the podocytes. Interestingly, in contrast to the paradigm that AMPK stimulates glycolysis under metabolically stressed conditions, we observed that elevated expression of essential glycolytic enzymes, such as PFKFB3 and PKM2, which promote glycolysis to generate pyruvate in the podocytes and other glomerular cells, such as endothelial cells, in diabetic pmut mice. These observations suggest that loss of AMPK in podocytes leads to a metabolic shift that is more preferential for utilizing glycolysis for fuel generation in glomerular cells under type 1 diabetic conditions. Furthermore, the isolated glomeruli from the pmut mice showed a higher level of LDH transcript whereas lower levels of PDH expression compared to control glomeruli under high glucose culture conditions, suggesting the tendency of production of lactate, which has been found to accelerate the fibrotic process in many cell types^92–97^. Interestingly, our observations suggest that loss of AMPK in podocytes accelerated mesenchymal activation (endothelial-to-mesenchymal transition, EndMT) in glomerular endothelial cells. The glomerular endothelial cells in the diabetic pmut mice also displayed more glycolytic enzymes with less CPT1a expression compared to those in control diabetic mice, suggestive of the metabolic shift in the endothelial cells. Under normal filtration conditions, glomerular endothelial cells provide the essential signals that maintain podocyte health and physiological functions. In contrast, hyperfiltration that occurs under diabetic conditions is thought to disrupt critical endothelial-podocyte crosstalk, resulting in the failure of glomerular functions^98–100^. It is postulated that podocytes also support endothelial cell health. Previous studies postulated that podocyte dysfunctions cause the increased permeability of the basement membrane, which may underlie the potential mechanisms whereby the metabolism of endothelial cells can be controlled by podocyte derangements under diabetic conditions ^101^. Further studies will be required to investigate the underlying mechanisms and factors that facilitate EndMT upon AMPK inhibition in the podocytes.

As outlined in Figure 8, while AMPK in podocytes is dispensable for physiological podocyte functions under the normal ad-lib fed condition, it plays an important role in maintaining the function of both podocyte and endothelial cells by reducing lipid toxicity and catalyzing appropriate levels of glucose to mitigate the induction of aberrant metabolism under diabetes. In the absence of AMPK in podocytes, pathological suppression of FAO and aberrant glycolysis are further exaggerated under type 1 diabetic conditions, leading to excessive lipid loads in the podocyte and neighboring endothelial cells, resulting in mitochondrial damage and activation of the c-GAS-STING pathway and inflammation. These accumulative effects of metabolic shift and inflammation may cause EndMT in the glomerular endothelial cells. While there are yet no known signals from podocytes in the AMPK-deficient state that may account for the proposed crosstalk between glomerular podocytes and endothelial cells, the present study proposes that AMPK activation would be a reliable approach to protect glomerular dysfunction from diabetes by maintaining the better metabolism in podocytes as well as other neighboring glomerular cells such as endothelial cells.

**Figure 8.**
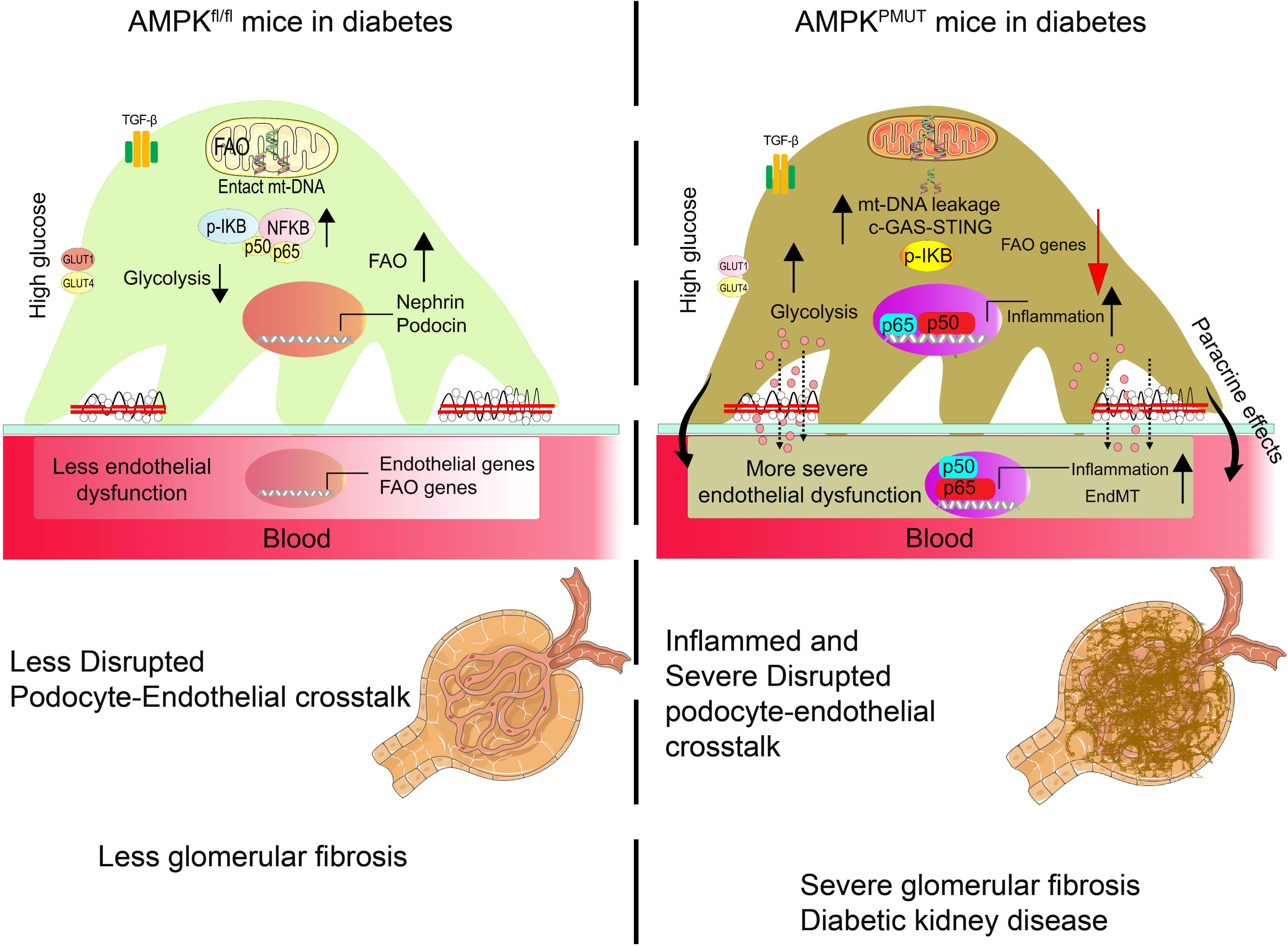
A hypothetical model of the proposed AMPK function in type 1 diabetic glomerular podocytes. While AMPK is dispensable in maintaining podocyte and glomerular function under normal fed conditions, with type 1 diabetes, AMPK may play an important role in mitigating the progression of DKD. Loss of AMPK in the podocytes exacerbated lipid deposition in the diabetic glomeruli, which in turn caused damage to mitochondria, leading to a leakage of mt-DNA into the cytoplasm and the activation of the cGAS-STING pathway. Activation of the STING pathway leads to the activation of p65 and enhances the transcription of genes associated with inflammation. Simultaneously, the loss of AMPK enhances aberrant glycolysis in the podocytes. These metabolic shifts in podocytes also instigate EndMT, and accumulative effects lead to the progression of glomerular fibrosis under diabetic conditions.

## Acknowledgments

We thank Junying Wang for her technical assistance in mice experiments and Mihir Bharadwaj for his help in image and statistical analyses.

## Author contributions

S.P.S. and K.I. designed, conceptualized, and performed experiments and wrote the paper. O.K-G., A.K., A.T., and E.O. also performed experiments. V.S. contributed to statistical analyses. S.Y. developed the image analysis program, and S.H. provided experimental methods.

---

**Supplemental Figure 1.**

**S1A.** Measurement of blood glucose in 2 years old WT and pmut mice using glucose strips. N=8 was analyzed/group.

**S1B.** Double immunofluorescence staining of pATG1 (S556) and WT1 was shown. While recent studies argue the role of phosphorylation of S566 of ATG1(ULK1) in regulating ATG1 activity, AMPK is a key kinase phosphorylating S556 residue. Representative images are shown. N=7/group were analyzed.

**S1C. Blood glucose level was determined in 2-year-old WT and pmut mice.** Data are shown as mean±SEM. N=8. ns indicates no significance.

**S1D.** Oil O Red staining was performed in the kidney tissues of 2-year-old WT and pmut mice. Representative images are shown.

## References

1 Jeon, S. M. Regulation and function of AMPK in physiology and diseases. Exp Mol Med 48, e245 (2016). 10.1038/emm.2016.81

2 Trefts, E. & Shaw, R. J. AMPK: restoring metabolic homeostasis over space and time. Mol Cell 81, 3677–3690 (2021). 10.1016/j.molcel.2021.08.015

3 Steinberg, G. R. & Hardie, D. G. New insights into activation and function of the AMPK. Nat Rev Mol Cell Biol 24, 255–272 (2023). 10.1038/s41580-022-00547-x

4 Oakhill, J. S. et al. AMPK is a direct adenylate charge-regulated protein kinase. Science 332, 1433–1435 (2011). 10.1126/science.1200094

5 Xiao, B. et al. Structure of mammalian AMPK and its regulation by ADP. Nature 472, 230–233 (2011). 10.1038/nature09932

6 Zhang, C. S. et al. Fructose-1,6-bisphosphate and aldolase mediate glucose sensing by AMPK. Nature 548, 112–116 (2017). 10.1038/nature23275

7 Zhang, Y. L. et al. AMP as a low-energy charge signal autonomously initiates assembly of AXIN-AMPK-LKB1 complex for AMPK activation. Cell Metab 18, 546–555 (2013). 10.1016/j.cmet.2013.09.005

8 Cooper, M. & Warren, A. M. A promising outlook for diabetic kidney disease. Nat Rev Nephrol 15, 68–70 (2019). 10.1038/s41581-018-0092-5

9 Gujarati, N. A. et al. Podocyte-specific KLF6 primes proximal tubule CaMK1D signaling to attenuate diabetic kidney disease. Nat Commun 15, 8038 (2024). 10.1038/s41467-024-52306-5

10 Zhao, Y. et al. Podocyte OTUD5 alleviates diabetic kidney disease through deubiquitinating TAK1 and reducing podocyte inflammation and injury. Nat Commun 15, 5441 (2024). 10.1038/s41467-024-49854-1

11 Thomas, M. C. et al. Diabetic kidney disease. Nat Rev Dis Primers 1, 15018 (2015). 10.1038/nrdp.2015.18

12 Tuttle, K. R. et al. Molecular mechanisms and therapeutic targets for diabetic kidney disease. Kidney Int 102, 248–260 (2022). 10.1016/j.kint.2022.05.012

13 Kim, M. K. Treatment of diabetic kidney disease: current and future targets. Korean J Intern Med 32, 622–630 (2017). 10.3904/kjim.2016.219

14 Dieter, B. P. et al. Novel Therapies for Diabetic Kidney Disease: Storied Past and Forward Paths. Diabetes Spectr 28, 167–174 (2015). 10.2337/diaspect.28.3.167

15 Pichler, R., Afkarian, M., Dieter, B. P. & Tuttle, K. R. Immunity and inflammation in diabetic kidney disease: translating mechanisms to biomarkers and treatment targets. Am J Physiol Renal Physiol 312, F716–F731 (2017). 10.1152/ajprenal.00314.2016

16 Bonnet, F., Cooper, M. E., Kopp, L., Fouque, D. & Candido, R. A review of the latest real-world evidence studies in diabetic kidney disease: What have we learned about clinical practice and the clinical effectiveness of interventions? Diabetes Obes Metab 26 Suppl 6, 55–65 (2024). 10.1111/dom.15710

17 Glastras, S. J. & Pollock, C. A. Targeted identification of risk and treatment of diabetic kidney disease. Nat Rev Nephrol 20, 75–76 (2024). 10.1038/s41581-023-00796-9

18 Kanasaki, K., Nangaku, M. & Ueki, K. ’DKD’ as the kidney disease relevant to individuals with diabetes. Diabetol Int 15, 673–676 (2024). 10.1007/s13340-024-00747-0

19 Kanasaki, K., Ueki, K. & Nangaku, M. Diabetic kidney disease: the kidney disease relevant to individuals with diabetes. Clin Exp Nephrol 28, 1213–1220 (2024). 10.1007/s10157-024-02537-z

20 Srivastava, S. P., Kanasaki, K. & Goodwin, J. E. Editorial: Combating Diabetes and Diabetic Kidney Disease. Front Pharmacol 12, 716029 (2021). 10.3389/fphar.2021.716029

21 Srivastava, S. P. & Kanasaki, K. Editorial: Receptor Biology and Cell Signaling in Diabetes. Front Pharmacol 13, 864117 (2022). 10.3389/fphar.2022.864117

22 Szrejder, M. et al. Metformin reduces TRPC6 expression through AMPK activation and modulates cytoskeleton dynamics in podocytes under diabetic conditions. Biochim Biophys Acta Mol Basis Dis 1866, 165610 (2020). 10.1016/j.bbadis.2019.165610

23 Lu, J. et al. GPR43 deficiency protects against podocyte insulin resistance in diabetic nephropathy through the restoration of AMPKalpha activity. Theranostics 11, 4728–4742 (2021). 10.7150/thno.56598

24 Li, X. et al. Inhibition of SGLT2 protects podocytes in diabetic kidney disease by rebalancing mitochondria-associated endoplasmic reticulum membranes. Cell Commun Signal 22, 534 (2024). 10.1186/s12964-024-01914-1

25 Lehtonen, S. Metformin Protects against Podocyte Injury in Diabetic Kidney Disease. Pharmaceuticals (Basel) 13 (2020). 10.3390/ph13120452

26 Han, Y. C. et al. AMPK agonist alleviate renal tubulointerstitial fibrosis via activating mitophagy in high fat and streptozotocin induced diabetic mice. Cell Death Dis 12, 925 (2021). 10.1038/s41419-021-04184-8

27 Salatto, C. T. et al. Selective Activation of AMPK beta1-Containing Isoforms Improves Kidney Function in a Rat Model of Diabetic Nephropathy. J Pharmacol Exp Ther 361, 303–311 (2017). 10.1124/jpet.116.237925

28 Zhou, X. et al. PAN-AMPK Activation Improves Renal Function in a Rat Model of Progressive Diabetic Nephropathy. J Pharmacol Exp Ther 371, 45–55 (2019). 10.1124/jpet.119.258244

29 Agur, T. et al. The impact of metformin on kidney disease progression and mortality in diabetic patients using SGLT2 inhibitors: a real-world cohort study. Cardiovasc Diabetol 24, 97 (2025). 10.1186/s12933-025-02643-6

30 Agranovich, A. L. et al. Carcinoid tumour of the gastrointestinal tract: prognostic factors and disease outcome. J Surg Oncol 47, 45–52 (1991). 10.1002/jso.2930470111

31 Srivastava, S. P. et al. Loss of endothelial glucocorticoid receptor accelerates diabetic nephropathy. Nat Commun 12, 2368 (2021). 10.1038/s41467-021-22617-y

32 Srivastava, S. P., Goodwin, J. E., Kanasaki, K. & Koya, D. Metabolic reprogramming by N-acetyl-seryl-aspartyl-lysyl-proline protects against diabetic kidney disease. Br J Pharmacol 177, 3691–3711 (2020). 10.1111/bph.15087

33 Srivastava, S. P., Goodwin, J. E., Kanasaki, K. & Koya, D. Inhibition of Angiotensin-Converting Enzyme Ameliorates Renal Fibrosis by Mitigating DPP-4 Level and Restoring Antifibrotic MicroRNAs. Genes (Basel) 11 (2020). 10.3390/genes11020211

34 Kanasaki, K. et al. Linagliptin-mediated DPP-4 inhibition ameliorates kidney fibrosis in streptozotocin-induced diabetic mice by inhibiting endothelial-to-mesenchymal transition in a therapeutic regimen. Diabetes 63, 2120–2131 (2014). 10.2337/db13-1029

35 Wang, H. et al. A simple and highly purified method for isolation of glomeruli from the mouse kidney. Am J Physiol Renal Physiol 317, F1217–F1223 (2019). 10.1152/ajprenal.00293.2019

36 Park, J. M., Lee, D. H. & Kim, D. H. Redefining the role of AMPK in autophagy and the energy stress response. Nat Commun 14, 2994 (2023). 10.1038/s41467-023-38401-z

37 Jia, J. et al. AMPK, a Regulator of Metabolism and Autophagy, Is Activated by Lysosomal Damage via a Novel Galectin-Directed Ubiquitin Signal Transduction System. Mol Cell 77, 951–969 e959 (2020). 10.1016/j.molcel.2019.12.028

38 Hartleben, B. et al. Autophagy influences glomerular disease susceptibility and maintains podocyte homeostasis in aging mice. The Journal of clinical investigation 120, 1084–1096 (2010). 10.1172/JCI39492

39 Srivastava, S. P. et al. Endothelial SIRT3 regulates myofibroblast metabolic shifts in diabetic kidneys. iScience 24, 102390 (2021). 10.1016/j.isci.2021.102390

40 Kang, H. M. et al. Defective fatty acid oxidation in renal tubular epithelial cells has a key role in kidney fibrosis development. Nat Med 21, 37–46 (2015). 10.1038/nm.3762

41 Zhou, H. L. et al. Metabolic reprogramming by the S-nitroso-CoA reductase system protects against kidney injury. Nature 565, 96–100 (2019). 10.1038/s41586-018-0749-z

42 Schoors, S. et al. Fatty acid carbon is essential for dNTP synthesis in endothelial cells. Nature 520, 192–197 (2015). 10.1038/nature14362

43 Tian, Z. & Liang, M. Renal metabolism and hypertension. Nat Commun 12, 963 (2021). 10.1038/s41467-021-21301-5

44 Huang, H. & Parikh, S. M. Skimming the fat in diabetic kidney disease: KIM-1 and tubular fatty acid uptake. Kidney Int 100, 969–972 (2021). 10.1016/j.kint.2021.06.038

45 Ralto, K. M., Rhee, E. P. & Parikh, S. M. NAD(+) homeostasis in renal health and disease. Nat Rev Nephrol 16, 99–111 (2020). 10.1038/s41581-019-0216-6

46 Tran, M. T. et al. PGC1alpha drives NAD biosynthesis linking oxidative metabolism to renal protection. Nature 531, 528–532 (2016). 10.1038/nature17184

47 Srivastava, S. P., Kanasaki, K. & Goodwin, J. E. Loss of Mitochondrial Control Impacts Renal Health. Front Pharmacol 11, 543973 (2020). 10.3389/fphar.2020.543973

48 Srivastava, S. P., Shi, S., Koya, D. & Kanasaki, K. Lipid mediators in diabetic nephropathy. Fibrogenesis Tissue Repair 7, 12 (2014). 10.1186/1755-1536-7-12

49 Srivastava, S. P. & Goodwin, J. E. Loss of endothelial glucocorticoid receptor accelerates organ fibrosis in db/db mice. Am J Physiol Renal Physiol (2023). 10.1152/ajprenal.00105.2023

50 Li, J. et al. Endothelial FGFR1 (Fibroblast Growth Factor Receptor 1) Deficiency Contributes Differential Fibrogenic Effects in Kidney and Heart of Diabetic Mice. Hypertension 76, 1935–1944 (2020). 10.1161/HYPERTENSIONAHA.120.15587

51 Shi, S. et al. Interactions of DPP-4 and integrin beta1 influences endothelial-to-mesenchymal transition. Kidney Int 88, 479–489 (2015). 10.1038/ki.2015.103

52 Zhou, H. et al. Endothelial cell-glucocorticoid receptor interactions and regulation of Wnt signaling. JCI Insight 5 (2020). 10.1172/jci.insight.131384

53 Srivastava, S. P., Hedayat, A. F., Kanasaki, K. & Goodwin, J. E. microRNA Crosstalk Influences Epithelial-to-Mesenchymal, Endothelial-to-Mesenchymal, and Macrophage-to-Mesenchymal Transitions in the Kidney. Front Pharmacol 10, 904 (2019). 10.3389/fphar.2019.00904

54 Li, J. et al. FGFR1 is critical for the anti-endothelial mesenchymal transition effect of N-acetyl-seryl-aspartyl-lysyl-proline via induction of the MAP4K4 pathway. Cell Death Dis 8, e2965 (2017). 10.1038/cddis.2017.353

55 Srivastava, S. P., Koya, D. & Kanasaki, K. MicroRNAs in kidney fibrosis and diabetic nephropathy: roles on EMT and EndMT. Biomed Res Int 2013, 125469 (2013). 10.1155/2013/125469

56 Srivastava, S. P., Goodwin, J. E., Tripathi, P., Kanasaki, K. & Koya, D. Interactions among Long Non-Coding RNAs and microRNAs Influence Disease Phenotype in Diabetes and Diabetic Kidney Disease. Int J Mol Sci 22 (2021). 10.3390/ijms22116027

57 Lovisa, S. & Kalluri, R. Fatty Acid Oxidation Regulates the Activation of Endothelial-to-Mesenchymal Transition. Trends Mol Med 24, 432–434 (2018). 10.1016/j.molmed.2018.03.003

58 Srivastava, S. P. et al. Renal Angptl4 is a key fibrogenic molecule in progressive diabetic kidney disease. Sci Adv 10, eadn6068 (2024). 10.1126/sciadv.adn6068

59 Wang, X. et al. Renal lipid accumulation and aging linked to tubular cells injury via ANGPTL4. Mech Ageing Dev 219, 111932 (2024). 10.1016/j.mad.2024.111932

60 Chung, K. W. et al. Mitochondrial Damage and Activation of the STING Pathway Lead to Renal Inflammation and Fibrosis. Cell Metab 30, 784–799 e785 (2019). 10.1016/j.cmet.2019.08.003

61 Decout, A., Katz, J. D., Venkatraman, S. & Ablasser, A. The cGAS-STING pathway as a therapeutic target in inflammatory diseases. Nat Rev Immunol 21, 548–569 (2021). 10.1038/s41577-021-00524-z

62 Chen, M. et al. cGAS-STING pathway expression correlates with genomic instability and immune cell infiltration in breast cancer. NPJ Breast Cancer 10, 1 (2024). 10.1038/s41523-023-00609-z

63 Nunes, J. R. C. et al. Myeloid AMPK signaling restricts fibrosis but is not required for metformin improvements during CDAHFD-induced NASH in mice. J Lipid Res 65, 100564 (2024). 10.1016/j.jlr.2024.100564

64 Wei, X. et al. Ulinastatin attenuates renal fibrosis by regulating AMPK/HIF-1alpha signaling pathway-mediated glycolysis. Sci Rep 14, 28032 (2024). 10.1038/s41598-024-78092-0

65 Wang, Y. et al. AMP-activated protein kinase/myocardin-related transcription factor-A signaling regulates fibroblast activation and renal fibrosis. Kidney Int 93, 81–94 (2018). 10.1016/j.kint.2017.04.033

66 Yanagi, T. et al. ULK1-regulated AMP sensing by AMPK and its application for the treatment of chronic kidney disease. Kidney Int 106, 887–906 (2024). 10.1016/j.kint.2024.08.024

67 Kanasaki, M. et al. Deficiency in catechol-o-methyltransferase is linked to a disruption of glucose homeostasis in mice. Sci Rep 7, 7927 (2017). 10.1038/s41598-017-08513-w

68 Lee, J. Y. et al. Ipragliflozin, an SGLT2 Inhibitor, Ameliorates High-Fat Diet-Induced Metabolic Changes by Upregulating Energy Expenditure through Activation of the AMPK/ SIRT1 Pathway. Diabetes Metab J 45, 921–932 (2021). 10.4093/dmj.2020.0187

69 Koh, E. S. et al. Anthocyanin-rich Seoritae extract ameliorates renal lipotoxicity via activation of AMP-activated protein kinase in diabetic mice. J Transl Med 13, 203 (2015). 10.1186/s12967-015-0563-4

70 Ye, K., Zhao, Y., Huang, W. & Zhu, Y. Sodium butyrate improves renal injury in diabetic nephropathy through AMPK/SIRT1/PGC-1alpha signaling pathway. Sci Rep 14, 17867 (2024). 10.1038/s41598-024-68227-8

71 Zhang, C. et al. Morroniside Ameliorates High-Fat and High-Fructose-Driven Chronic Kidney Disease by Motivating AMPK-TFEB Signal Activation to Accelerate Lipophagy and Inhibiting Inflammatory Response. J Agric Food Chem 73, 6158–6172 (2025). 10.1021/acs.jafc.4c07684

72 Liu, X. et al. Empagliflozin improves diabetic renal tubular injury by alleviating mitochondrial fission via AMPK/SP1/PGAM5 pathway. Metabolism 111, 154334 (2020). 10.1016/j.metabol.2020.154334

73 Kim, M. Y. et al. Resveratrol prevents renal lipotoxicity and inhibits mesangial cell glucotoxicity in a manner dependent on the AMPK-SIRT1-PGC1alpha axis in db/db mice. Diabetologia 56, 204–217 (2013). 10.1007/s00125-012-2747-2

74 Hong, Y. A. et al. Fenofibrate improves renal lipotoxicity through activation of AMPK-PGC-1alpha in db/db mice. PLoS One 9, e96147 (2014). 10.1371/journal.pone.0096147

75 Godel, M. et al. Role of mTOR in podocyte function and diabetic nephropathy in humans and mice. J Clin Invest 121, 2197–2209 (2011). 10.1172/JCI44774

76 Inoki, K. et al. mTORC1 activation in podocytes is a critical step in the development of diabetic nephropathy in mice. J Clin Invest 121, 2181–2196 (2011). 10.1172/JCI44771

77 Puelles, V. G. et al. mTOR-mediated podocyte hypertrophy regulates glomerular integrity in mice and humans. JCI Insight 4 (2019). 10.1172/jci.insight.99271

78 Inoki, K. et al. TSC2 integrates Wnt and energy signals via a coordinated phosphorylation by AMPK and GSK3 to regulate cell growth. Cell 126, 955–968 (2006). 10.1016/j.cell.2006.06.055

79 Centers for Disease, C. Progress toward achieving the 1990 high blood pressure objectives. MMWR Morb Mortal Wkly Rep 39, 704–707 (1990).

80 Kim, J., Kundu, M., Viollet, B. & Guan, K. L. AMPK and mTOR regulate autophagy through direct phosphorylation of Ulk1. Nat Cell Biol 13, 132–141 (2011). 10.1038/ncb2152

81 Iwata, W. et al. Podocyte-specific deletion of tubular sclerosis complex 2 promotes focal segmental glomerulosclerosis and progressive renal failure. PLoS One 15, e0229397 (2020). 10.1371/journal.pone.0229397

82 Bork, T. et al. Podocytes maintain high basal levels of autophagy independent of mtor signaling. Autophagy 16, 1932–1948 (2020). 10.1080/15548627.2019.1705007

83 Dugan, L. L. et al. AMPK dysregulation promotes diabetes-related reduction of superoxide and mitochondrial function. J Clin Invest 123, 4888–4899 (2013). 10.1172/JCI66218

84 Banu, K. et al. AMPK mediates regulation of glomerular volume and podocyte survival. JCI Insight 6 (2021). 10.1172/jci.insight.150004

85 Formigari, G. P. et al. Renal protection induced by physical exercise may be mediated by the irisin/AMPK axis in diabetic nephropathy. Sci Rep 12, 9062 (2022). 10.1038/s41598-022-13054-y

86 Lee, H. J. et al. Female Protection Against Diabetic Kidney Disease Is Regulated by Kidney-Specific AMPK Activity. Diabetes 73, 1167–1177 (2024). 10.2337/db23-0807

87 Decleves A-E, Zolkipli Z, Satriano J, et al. Regulation of lipid accumulation by AMK-activated kinase in high fat diet-induced kidney injury. Kidney Int. 2014;85:611-623. Kidney Int 92, 769 (2017). 10.1016/j.kint.2017.06.011

88 Kim, D. et al. Metformin decreases high-fat diet-induced renal injury by regulating the expression of adipokines and the renal AMP-activated protein kinase/acetyl-CoA carboxylase pathway in mice. Int J Mol Med 32, 1293–1302 (2013). 10.3892/ijmm.2013.1508

89 Rodriguez, C. et al. Activation of AMP kinase ameliorates kidney vascular dysfunction, oxidative stress and inflammation in rodent models of obesity. Br J Pharmacol 178, 4085–4103 (2021). 10.1111/bph.15600

90 Lee, H. J. et al. Proximal tubular epithelial insulin receptor mediates high-fat diet-induced kidney injury. JCI Insight 6 (2021). 10.1172/jci.insight.143619

91 Wu, L. et al. The Attenuation of Diabetic Nephropathy by Annexin A1 via Regulation of Lipid Metabolism Through the AMPK/PPARalpha/CPT1b Pathway. Diabetes 70, 2192–2203 (2021). 10.2337/db21-0050

92 Srivastava, S. P. et al. SIRT3 deficiency leads to induction of abnormal glycolysis in diabetic kidney with fibrosis. Cell Death Dis 9, 997 (2018). 10.1038/s41419-018-1057-0

93 Li, J. et al. Renal protective effects of empagliflozin via inhibition of EMT and aberrant glycolysis in proximal tubules. JCI Insight 5 (2020). 10.1172/jci.insight.129034

94 Wang, Y. et al. The glycolytic enzyme PFKFB3 drives kidney fibrosis through promoting histone lactylation-mediated NF-kappaB family activation. Kidney Int 106, 226–240 (2024). 10.1016/j.kint.2024.04.016

95 Zhao, B. et al. Activation of TRPV4 by lactate as a critical mediator of renal fibrosis in spontaneously hypertensive rats after moderate- and high-intensity exercise. Front Physiol 13, 927078 (2022). 10.3389/fphys.2022.927078

96 Zhang, X. et al. Lactate drives epithelial-mesenchymal transition in diabetic kidney disease via the H3K14la/KLF5 pathway. Redox Biol 75, 103246 (2024). 10.1016/j.redox.2024.103246

97 Shen, Y. et al. Tubule-derived lactate is required for fibroblast activation in acute kidney injury. Am J Physiol Renal Physiol 318, F689–F701 (2020). 10.1152/ajprenal.00229.2019

98 Hu, S. et al. Crosstalk among podocytes, glomerular endothelial cells and mesangial cells in diabetic kidney disease: an updated review. Cell Commun Signal 22, 136 (2024). 10.1186/s12964-024-01502-3

99 Clark, A. R. et al. Single-Cell Transcriptomics Reveal Disrupted Kidney Filter Cell-Cell Interactions after Early and Selective Podocyte Injury. Am J Pathol 192, 281–294 (2022). 10.1016/j.ajpath.2021.11.004

100 Greka, A. & Mundel, P. Cell biology and pathology of podocytes. Annu Rev Physiol 74, 299–323 (2012). 10.1146/annurev-physiol-020911-153238

101 Srivastava, S. P. et al. Podocyte Glucocorticoid Receptors Are Essential for Glomerular Endothelial Cell Homeostasis in Diabetes Mellitus. J Am Heart Assoc 10, e019437 (2021). 10.1161/JAHA.120.019437

