## Supplementary figures and images for "AMPK is dispensable for physiological podocyte and glomerular functions but prevents glomerular fibrosis in experimental diabetes"

### supplemental figure 1

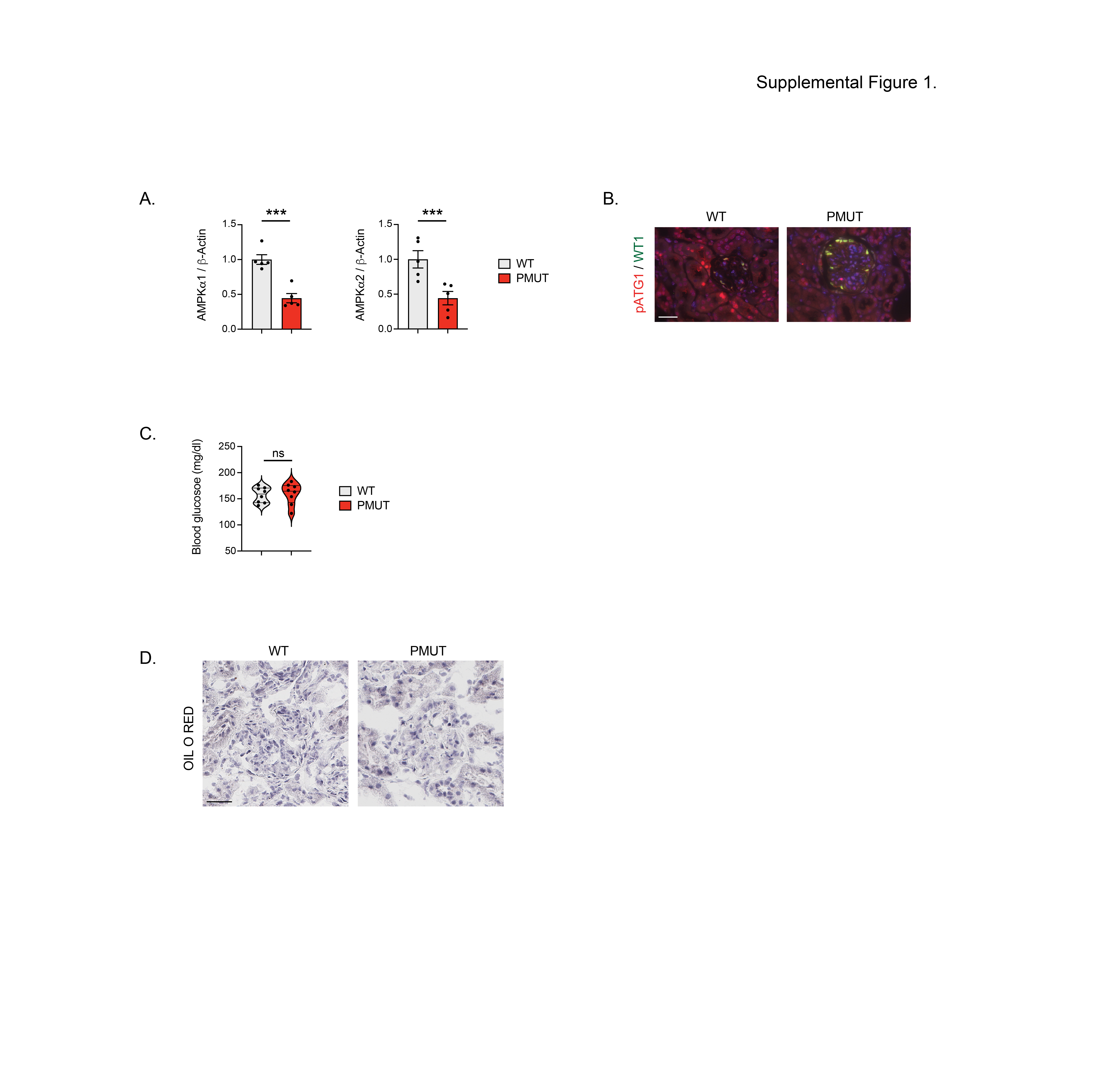
