## Supplementary table 1 for "AMPK is dispensable for physiological podocyte and glomerular functions but prevents glomerular fibrosis in experimental diabetes"

**Table S1: Sequences mouse primers used**

| Gene name | Forward primer | Reverse primer |
| --- | --- | --- |
| NFkB1 | 5′-GCTGCCAAAGAAGGACACGACA | 5′-GGCAGGCTATTGCTCATCACAG |
| IL-1β | 5′-TGGACCTTCCAGGATGAGGACA | 5′-GTTCATCTCGGAGCCTGTAGTG |
| IL-6 | 5′-TCTGAAGGACTCTGGCTTTG | 5′-GATGGATGCTACCAAACTGGA |
| Ndufv2 | 5’-TGGATGGCTACCTATCTCCGCT | 5’-GGTACTTCCCAACTGGCTTTCG |
| PDHA1 | 5′-GTGAGAACAACCGCTATGGCATG | 5′-CGCAAACTTTGTTGCCTCTCGG |
| LDH | 5’-ACGCAGACAAGGAGCAGTGGAA | 5’-ATGCTCTCAGCCAAGTCTGCCA |
| mt-Co1 | 5’-GCCCCAGATATAGCATTCCC | 5’-GTTCATCCTGTTCCTGCTCC |
| mt-Cyb | 5’-AGTAGACAAAGCCACCTTGA | 5’-CCGCGATAATAAATGGTAAG |
| TFAM | 5’-GAGCAGCTAACTCCAAGTCAG | 5’-GAGCCGAATCATCCTTTGCCT |
| TMEM173 | 5’-TTTGCCATGTCACAGGATGC | 5’-ATGAGGCGGCAGTTATTTCG |
| Mb21d1 | 5’-TGGTGGGAAGAGTGGTGATTTC | 5’-TGCATTCCAATGGCAGAAGC |
| β-actin | 5’-AGAAGCTGTGCTATGTTGCTCTA | 5’-ACAGGATTCCATACCCAAGAAGGA |
